# Gram-negative-dominated polymicrobial microbiome of necrotizing soft tissue infections from North India: an integrated culture and 16S rRNA metagenomics prospective cohort study

**DOI:** 10.64898/2026.06.25.734553

**Authors:** Tanvi Vashist, Nitika Rana, Divya Nair, Vikas Sharma, Anjali Anil, Cherring Tandup, Pallab Ray, Archana Angrup

## Abstract

Necrotizing soft tissue infections (NSTIs) carry 10 to 30% mortality. Current empirical antimicrobial guidance derives almost entirely from Western cohorts dominated by *Streptococcus pyogenes* and aerobic-anaerobic consortia, yet whether this microbial paradigm applies to tropical, high-antimicrobial-pressure settings has not been tested with culture-independent methods. We did a prospective cohort study of 169 patients with intraoperatively confirmed NSTI at a North Indian tertiary center (2021 to 2024). Wound tissue underwent aerobic and anaerobic culture, QIIME2-based 16S rRNA gene amplicon sequencing (V3-V4), and targeted SYBR Green quantitative PCR (qPCR) for *Acinetobacter baumannii* and *S. pyogenes*. The wound microbiota was overwhelmingly Gram-negative and polymicrobial, anchored by *A. baumannii* (culture, 33.7%; metagenomics, 49.1%; qPCR, 37.9%), *Escherichia coli* (32.0%), and *Klebsiella pneumoniae* (20.7%); *S. pyogenes* contributed only 4.7% of culture-positive cases. Polymicrobial wounds had higher Shannon diversity (2.59 versus 2.33; *P* = 0.048) and discrete community composition (PERMANOVA *R*^2^ = 0.511; *P* = 0.010). Culture-metagenomics agreement ranged from almost perfect for *Escherichia* (κ = 0.849) to slight for *Streptococcus* (κ = 0.131). North Indian NSTIs present a microbial picture distinct from the Western paradigm, with implications for empirical therapy.

**IMPORTANCE:** Necrotizing soft tissue infections kill rapidly, and physicians must choose antibiotics before laboratory results are available. Globally, treatment guidelines assume the dominant pathogen is *Streptococcus pyogenes*, a Gram-positive organism, because nearly all molecular evidence comes from Western countries. We show that in a large Indian cohort, the infection is instead dominated by Gram-negative bacteria, especially *Acinetobacter baumannii*, with *S. pyogenes* rarely present. Polymicrobial status, not wound location or injury mechanism, most strongly determines community composition. These findings challenge the one-size-fits-all antibiotic approach and argue for region-specific empirical regimens prioritizing Gram-negative coverage in tropical, high-antimicrobial-pressure settings.

## INTRODUCTION

Necrotizing soft tissue infections (NSTIs) are rapidly progressive infections of the subcutaneous fascia and muscle that carry a mortality of 10 to 30% despite prompt surgical debridement and broad-spectrum antimicrobial therapy (1, 2). Outcome depends critically on the speed of source control and on the appropriateness of empirical antibiotic cover initiated before microbiological confirmation. The historical microbiological framework distinguishes type I disease (polymicrobial, aerobic-anaerobic) from type II disease (monomicrobial, predominantly *Streptococcus pyogenes*) (1, 3, 4) and underpins current empirical antimicrobial guidance worldwide. This taxonomy, and the antibiotic regimens built upon it, were constructed almost entirely from Western, temperate-climate cohorts.

The most rigorous molecular characterization of NSTI to date is the multicenter INFECT study, which combined 16S rRNA gene amplicon sequencing with dual host-pathogen RNA-seq in 148 Scandinavian patients and reaffirmed *S. pyogenes* dominance in monomicrobial disease alongside anatomically structured anaerobic communities in polymicrobial infection (5). This single dataset, together with smaller Western series, anchors current understanding of the NSTI wound microbiome. The broader wound-microbiome literature, however, indicates that the microbial composition of soft tissue infection is shaped by host comorbidity, environmental exposure, and prior antimicrobial pressure (6, 7), and that polymicrobial community structure rather than any single isolate often predicts clinical trajectory (8). Whether the INFECT pattern extends to populations outside northern Europe is therefore an empirical question, not a settled one.

Indian and broader South Asian case series consistently report aerobic Gram-negative organisms, particularly *Acinetobacter baumannii*, *Escherichia coli*, and *Klebsiella pneumoniae*, as the predominant culture isolates from necrotizing soft tissue disease (9, 10). *A. baumannii* is of particular clinical concern: it is environmentally persistent, frequently multidrug resistant, biofilm forming, and has emerging recognition as a primary pathogen in necrotizing fasciitis (11–13). Yet, to our knowledge, no published study had applied culture-independent 16S metagenomics to NSTI in an Indian cohort, and the diagnostic performance of culture, quantitative PCR (qPCR), and amplicon sequencing in tropical NSTI had not been compared in a chance-corrected framework. This evidence gap directly affects empirical antimicrobial choice in the highest-mortality phase of clinical care.

We therefore did an integrated, multimodal characterization of the NSTI wound microbiome from the Indian subcontinent. In a prospective cohort of 169 patients at a North Indian tertiary referral center, we combined aerobic and anaerobic culture, QIIME2-based 16S rRNA gene amplicon sequencing, and targeted SYBR Green qPCR to address three aims: (i) identify the organisms that dominate North Indian NSTIs and compare this with the Western paradigm, (ii) determine which clinical variable, anatomic site, traumatic etiology, or polymicrobial status, most strongly shapes the wound microbial community, and (iii) compare culture, qPCR, and amplicon sequencing as diagnostic modalities using both one-directional sensitivity and chance-corrected Cohen’s κ.

## MATERIALS AND METHODS

### Study design and patients

This prospective observational cohort study enrolled patients with intraoperatively confirmed NSTIs at the Postgraduate Institute of Medical Education and Research (PGIMER), Chandigarh, North India, between 2021 and 2024. The Institutional Ethics Committee approved the study (IEC-INT/2022/PhD-406), and the work was done in accordance with the Declaration of Helsinki; written informed consent was obtained from all participants. NSTI diagnosis required intraoperative confirmation of necrotizing infection involving fascia or muscle as determined by the operating surgeon. Of 270 patients evaluated, 169 met the inclusion criteria after excluding those with insufficient sample material, those who were pregnant, or those who declined consent (see Fig. S1 in the supplemental material). Patients were stratified by anatomic site (group A, extremities, *n* = 126; group B, nonextremity, *n* = 43), traumatic etiology (traumatic, *n* = 68; nontraumatic, *n* = 101), and polymicrobial status, defined by culture as at least two distinct pathogenic species recovered from a single biopsy. Sample size was determined by feasibility; consecutive eligible patients were enrolled over three years, and no formal power calculation was done. The study is reported in accordance with the STROBE statement (see Table S1 in the supplemental material).

### Microbiological culture

Intraoperative tissue biopsies were inoculated onto sheep blood agar and MacConkey agar (aerobic incubation, 37°C, 24 to 48 h) and anaerobic blood agar in GasPak chambers (48 to 72 h). Bacterial isolates were identified by matrix-assisted laser desorption ionization–time of flight (MALDI-TOF) mass spectrometry (Vitek MS; bioMérieux, France).

### 16S rRNA gene amplicon sequencing

Genomic DNA was extracted from RNAlater-preserved biopsies using the DNeasy blood and tissue kit (Qiagen, Germany) and quantified by NanoDrop and Qubit fluorometry. A universal 16S rRNA PCR screen (primers DG74/RW01, 379 bp) excluded 34 samples without amplifiable bacterial DNA. The hypervariable V3-V4 region was amplified using primers 357F/805R according to the Illumina 16S metagenomic library preparation protocol and sequenced on a MiSeq instrument (2 × 300 bp). Three sequencing runs were batch corrected using MBECS v1.4.0 in R v4.3.1. Extraction blanks and PCR no-template controls were not included in this sequencing run; this limitation is discussed below.

### Bioinformatic processing

Raw paired-end reads underwent quality assessment with FastQC v0.12.1 (29) and adapter trimming with Trimmomatic v0.39 (20); aggregate quality control was generated with MultiQC (21). Reads were imported into QIIME2 v2023.2 (16) and processed with the DADA2 plugin (17) using truncation lengths of 275 bp (forward) and 260 bp (reverse) determined from per-base quality profiles. DADA2 performed phiX read removal, error-model-based denoising, paired-end merging, and *de novo* chimera filtering. DADA2 processing yielded 182 ASVs from 3102 raw input sequences; after low-count filtering, 170 genus-level features were retained for downstream analysis. Counts were normalized by total-sum scaling. Taxonomy was assigned using a naive Bayes classifier trained against the SILVA 138 reference database within QIIME2; features were collapsed to genus level for all downstream analyses. The pipeline is summarized in the supplemental material.

### Statistical analysis

Alpha diversity (Shannon *H*′, Simpson 1 − *D*, Chao1 estimated richness, observed amplicon sequence variants [ASVs]) was computed from the rarefied feature table at a rarefaction depth of 900 reads per sample. Continuous variables were not assumed to be normally distributed; group differences were tested by the two-sided Mann-Whitney *U* test, with Benjamini-Hochberg false-discovery-rate (FDR) correction applied across all multiple comparisons. Where parametric comparison was appropriate (after visual and Shapiro-Wilk assessment), Welch’s *t* test or one-way analysis of variance (ANOVA) was used. Beta diversity was assessed using Bray-Curtis dissimilarity and visualized by principal-coordinate analysis (PCoA); group differences were tested by permutational multivariate analysis of variance (PERMANOVA) with 999 permutations. Homogeneity of multivariate group dispersions was tested by PERMDISP (30) on the first five PCoA axes (18.5% cumulative variance) with 999 permutations. Differential genus abundance between polymicrobial and monomicrobial communities was assessed by linear discriminant analysis effect size LEfSe; linear discriminant analysis with a LDA score of ≥2.0 used as the primary discriminant criterion. Co-occurrence networks were inferred from genus-level relative abundances using Spearman rank correlation with SparCC permutations (*n* = 100); only edges with absolute correlation ≥0.3 and adjusted *P* ≤ 0.05 were retained. Culture-metagenomics agreement was quantified by both one-directional sensitivity (true positives/[true positives + false negatives]) and chance-corrected Cohen’s κ with Landis-Koch interpretation bands; the Matthews correlation coefficient was computed as a complementary, class-imbalance-robust measure (see the supplemental material). Spearman rank correlation related qPCR cycle threshold (*C*_T_) values to metagenomic relative abundance. The alpha level was 0.05 throughout, with two-tailed testing unless otherwise specified. All analyses were done in Python v3.10 (SciPy v1.11, NumPy v1.24, and pandas v2.0).

### qPCR validation

Targeted SYBR Green qPCR was performed for *A. baumannii* (*omp* gene; all 169 samples) and for *S. pyogenes* (M1 protein; 139 samples with sufficient residual DNA). Primer sequences are listed in the supplemental material. Primer sequences and confirmed producte sizes are listed in Table S1 in the supplemental materials. Reactions (10 μL) contained 5 μL of 2× SYBR master mix, 0.5 μM each primer, and 2 μL of template DNA. Cycling conditions were 95°C for 3 min, followed by 40 cycles of 95°C for 30 s, 54°C for 30 s, and 72°C for 30 s. Specificity was confirmed by melt-curve analysis from 65°C to 95°C. A *C*_T_ value of ≤35 was considered positive.

## RESULTS

### Patient cohort

A total of 169 patients (138 [81.7%] male) were enrolled between 2021 and 2024; the mean age was 42.8 (standard deviation [SD], 16.2) years. Diabetes mellitus was the dominant comorbidity (52.1%). Group A (extremity infections) comprised 126 patients, and group B (nonextremity) comprised 43. A traumatic etiology was identified in 68 patients (40.2%). Culture classified 98 infections (58.0%) as polymicrobial, 65 (38.5%) as monomicrobial, and 6 (3.6%) as sterile; of the 98 polymicrobial cases, 97 yielded amplicon sequencing data (1 sample failed postfilter read-depth thresholds). Of the 98 culture-classified polymicrobial cases, 97 yielded sequencing data passing post-filter depth thresholds; the 6 sterile samples and remaining sequencing failures were excluded from the beta-diversity analyses (PERMANOVA n=163). Necrotizing fasciitis accounted for 55.0% of presentations. In-hospital mortality occurred in 28 of 147 patients with documented outcomes (19.05%; overall cohort mortality, 16.6%). Mortality outcomes were unavailable for 22 patients (13.0%) who were transferred or left against medical advice before discharge.

### Microbiological culture

Culture recovered 65 species (Table 1). *A. baumannii* was the most frequent isolate (57/169, 33.7%), followed by *E. coli* (54, 32.0%), *K. pneumoniae* (35, 20.7%), *Proteus mirabilis* (31, 18.3%), *Pseudomonas aeruginosa* (26, 15.4%), and *Staphylococcus aureus* (21, 12.4%). Anaerobes were infrequent (*Bacteroides ovatus*, 10, 5.9%). *S. pyogenes* was isolated in only 8 patients (4.7%), contrasting sharply with Western epidemiology. *A. baumannii* was strongly associated with polymicrobial infection (43/57, 75.4%) and extremity involvement (group A, 44; group B, 13).

**TABLE 1.**
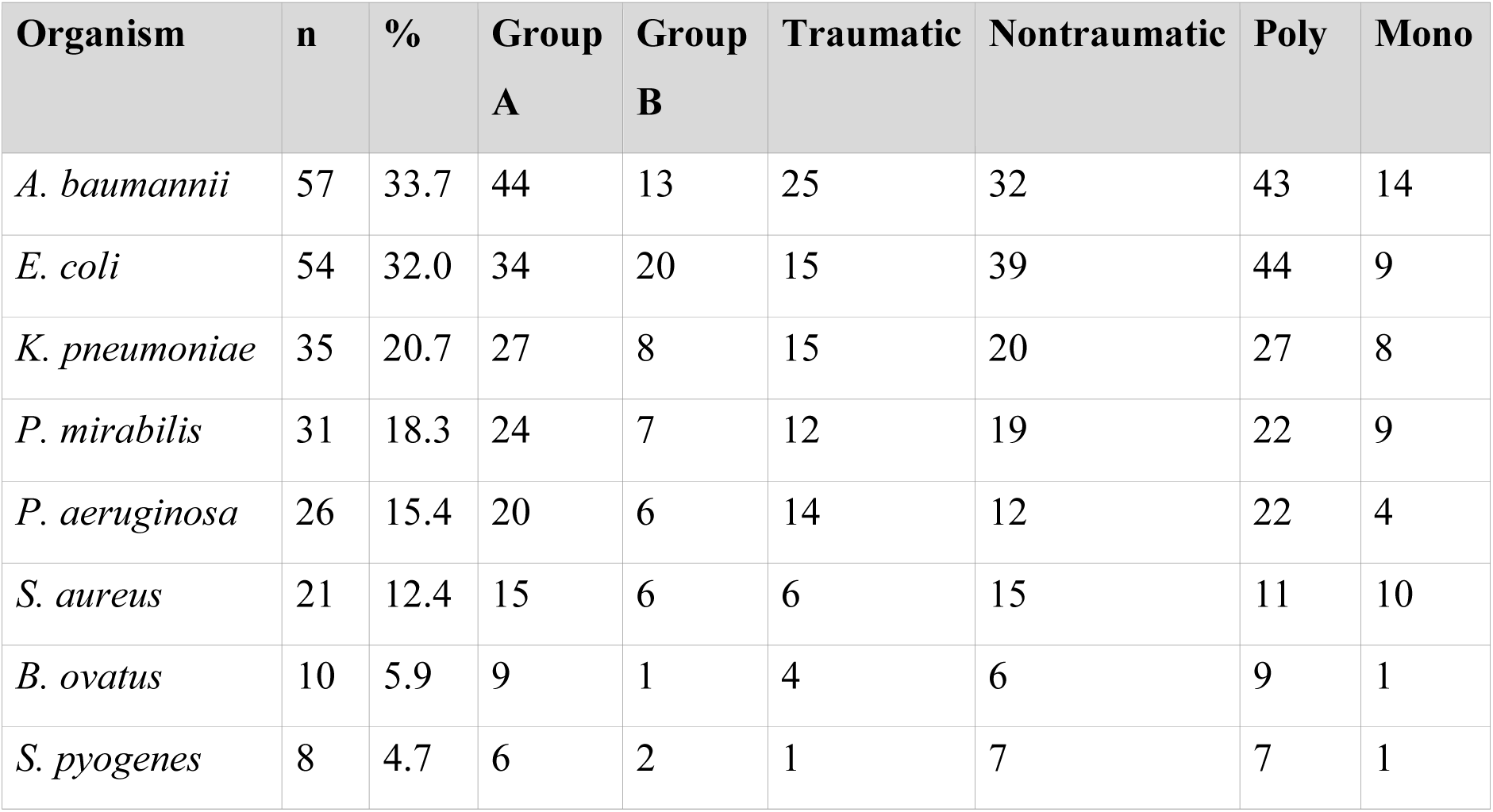
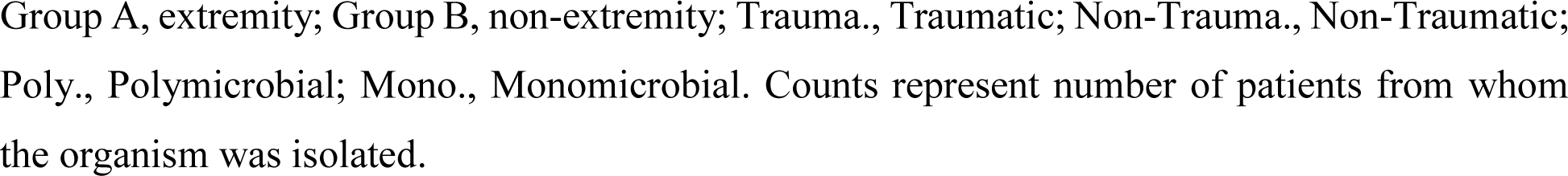
Predominant bacterial isolates from NSTI tissue biopsies (n = 169)

| Organism | n | % | Group<br>A | Group<br>B | Traumatic | Nontraumatic | Poly | Mono |
| --- | --- | --- | --- | --- | --- | --- | --- | --- |
| *A. baumannii* | 57 | 33.7 | 44 | 13 | 25 | 32 | 43 | 14 |
| *E. coli* | 54 | 32.0 | 34 | 20 | 15 | 39 | 44 | 9 |
| *K. pneumoniae* | 35 | 20.7 | 27 | 8 | 15 | 20 | 27 | 8 |
| *P. mirabilis* | 31 | 18.3 | 24 | 7 | 12 | 19 | 22 | 9 |
| *P. aeruginosa* | 26 | 15.4 | 20 | 6 | 14 | 12 | 22 | 4 |
| *S. aureus* | 21 | 12.4 | 15 | 6 | 6 | 15 | 11 | 10 |
| *B. ovatus* | 10 | 5.9 | 9 | 1 | 4 | 6 | 9 | 1 |
| *S. pyogenes* | 8 | 4.7 | 6 | 2 | 1 | 7 | 7 | 1 |

Group A, extremity; Group B, non-extremity; Trauma., Traumatic; Non-Trauma., Non-Traumatic; Poly., Polymicrobial; Mono., Monomicrobial. Counts represent number of patients from whom the organism was isolated.

### 16S metagenomics: taxonomic overview and core microbiome

Of the 169 enrolled patients, 167 yielded sequencing libraries that passed initial quality and contamination filters; 2 samples were excluded because of insufficient postfilter read depth. Sequencing returned 977,546 reads in total (mean, 5,818 per sample). After DADA2 denoising, chimera removal, and low-count filtering, 170 genus-level features were retained for downstream analysis. *Acinetobacter* was the dominant classified genus (mean relative abundance, 15.18%; sample prevalence, 49.1%), followed by *Escherichia* (7.44%; 31.7%), *Streptococcus* (6.74%; 30.5%), *Proteus* (5.86%; 19.2%), *Klebsiella* (3.53%; 18.0%), *Bacteroides* (3.25%; 24.6%), and *Staphylococcus* (2.32%; 26.9%). Unclassified taxa comprised 20.3% of reads (Fig. 1). Applying a core-microbiome threshold of at least 20% prevalence and at least 0.01% relative abundance, seven genera qualified as core members: *Acinetobacter*, *Escherichia*, *Streptococcus*, *Staphylococcus*, *Bacteroides*, *Anaerococcus*, and unclassified sequences. *Proteus* and *Klebsiella* fell marginally below this threshold (see the supplemental material).

**FIG 1.**
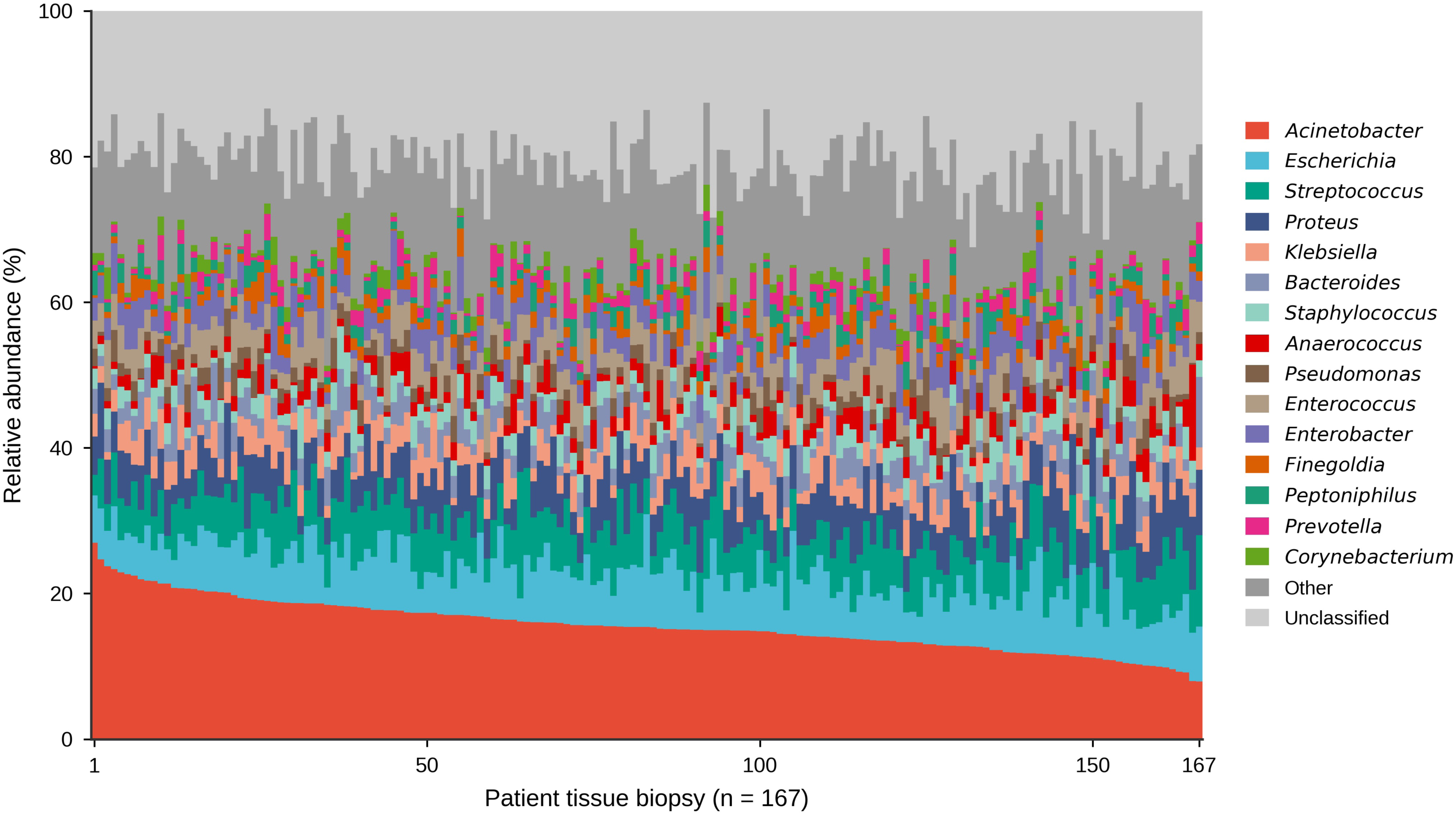
Genus-level taxonomic composition of the NSTI tissue microbiome across 167 patient samples. Each vertical bar represents a single patient tissue biopsy; colored stacked segments indicate the relative abundance of QIIME2-derived genera (V3-V4 16S rRNA amplicon sequencing; SILVA 138 taxonomy). “Unclassified” denotes sequences assigned only at the kingdom or phylum level.

### Alpha diversity

Shannon diversity ranged from 0 to 4.24 (mean, 2.50 [SD, 0.88]), Simpson from 0 to 0.98 (mean, 0.86 [0.16]), and observed ASV richness from 0 to 164 (median, 19 [interquartile range {IQR}, 9 to 39]). Polymicrobial NSTIs harbored significantly higher alpha diversity than monomicrobial infections: Shannon (2.59 [0.86] versus 2.33 [0.88]; *P* = 0.048), observed ASVs (median, 21 versus 13.5; *P* = 0.046), and Chao1 (*P* = 0.045). Simpson diversity showed a nonsignificant trend (0.87 [0.15] versus 0.83 [0.17]; *P* = 0.051). Neither anatomic site (Shannon *P* = 0.243) nor traumatic etiology (Shannon *P* = 0.391) had a measurable effect on alpha diversity (Table 2; Fig. 2). Benjamini-Hochberg FDR adjustment did not alter these conclusions.

**FIG 2.**
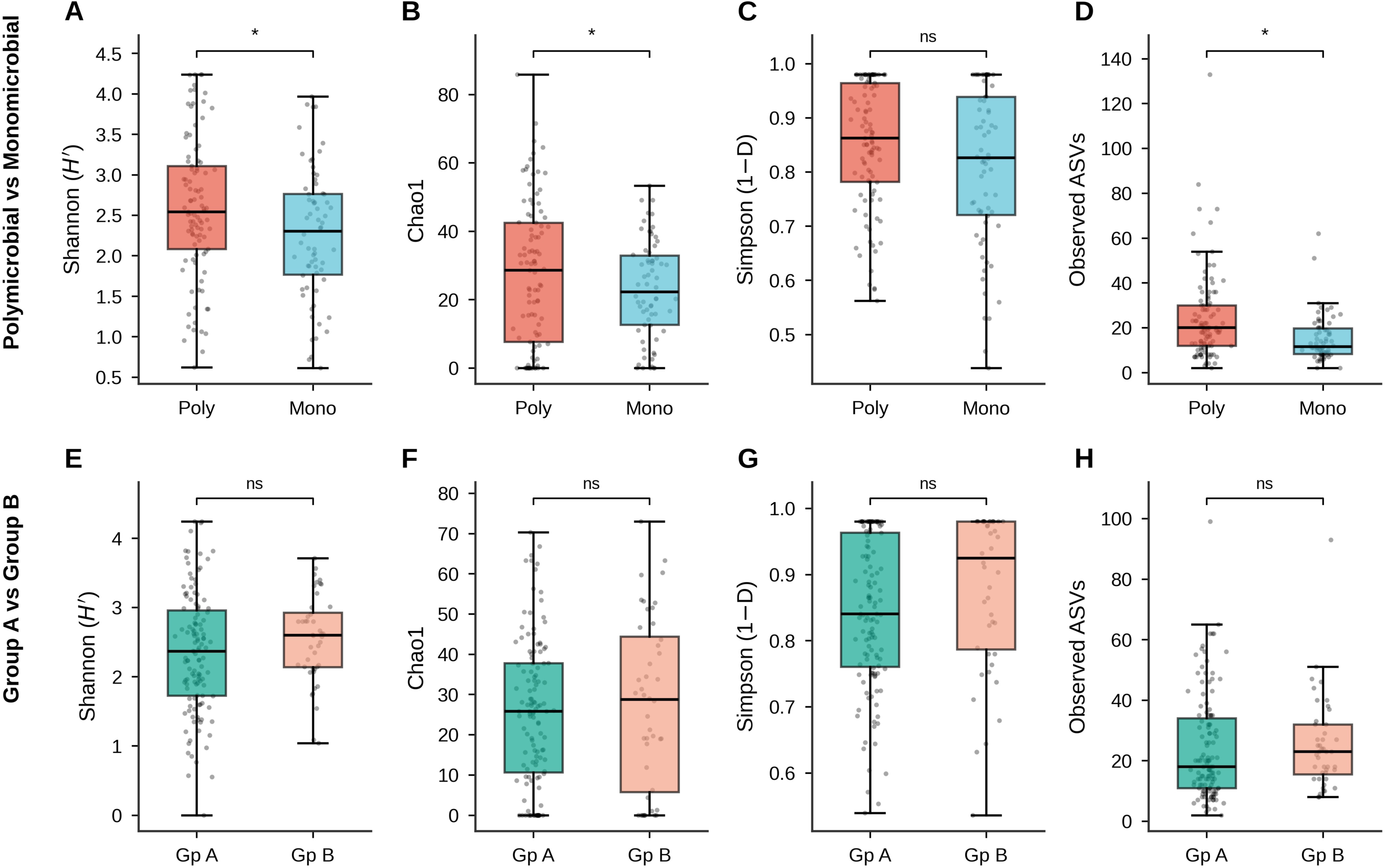
Alpha diversity across NSTI clinical subgroups. Box plots show four alpha diversity metrics (Shannon H′, Chao1, Simpson 1 − D, observed ASVs) for two comparisons: polymicrobial (n = 97) versus monomicrobial (n = 66) and group A (extremity; n = 123) versus group B (nonextremity; n = 44). Lines indicate medians; boxes span the IQR; whiskers extend to 1.5× IQR. Brackets show Mann-Whitney U P values after Benjamini-Hochberg FDR correction. ns, not significant. *, P < 0.05.

**Table 2.** Alpha diversity indices across NSTI clinical subgroups (mean ± s.d.).

| Subgroup (n) | Shanno<br>n | Simpson | Obs.<br>ASVs<br>(med.) | Chao1 | p-value |
| --- | --- | --- | --- | --- | --- |
| All (n = 167) | 2.50 ± 0.88 | 0.86 ± 0.16 | 19.0 | 24.6 ± 21.4 | , |
| Group A (n = 123) | 2.46 ± 0.84 | 0.86 ± 0.15 | 19.0 | 24.0 ± 21.1 | , |
| Group B (n = 44) | 2.57 ± 0.98 | 0.85 ± 0.18 | 19.5 | 26.8 ± 22.3 | , |
| p (A vs B) | 0.243 | 0.421 | 0.551 | 0.556 | ns |
| Traumatic (n = 67) | 2.55 ± 0.95 | 0.85 ± 0.19 | 21.0 | 25.9 ± 19.4 | , |
| Non-traumatic (n = 100) | 2.46 ± 0.83 | 0.86 ± 0.14 | 18.0 | 24.0 ± 22.6 | , |
| p (Traum. vs Non-traum.) | 0.391 | 0.462 | 0.406 | 0.421 | ns |
| Polymicrobial (n = 97) | 2.59 ± 0.86 | 0.87 ± 0.15 | 21.0 | 27.1 ± 23.4 | , |
| Monomicrobial (n = 66) | 2.33 ± 0.88 | 0.83 ± 0.17 | 13.5 | 20.6 ± 17.2 | , |
| p (Poly vs Mono) | 0.048* | 0.051 | 0.046* | 0.045* | * |

Mann-Whitney U test (two-sided) with Benjamini-Hochberg FDR correction. \* $p < 0.05$ ; ns, not significant.

### Beta diversity and community composition

Bray-Curtis PCoA axes 1 and 2 explained 6.19% and 4.34% of total variance, respectively (cumulative, 10.53%), consistent with substantial interindividual heterogeneity. PERMANOVA revealed significant compositional divergence between polymicrobial and monomicrobial NSTIs (pseudo-*F* = 85.55; *R*^2^ = 0.511; *P* = 0.010), with polymicrobial status accounting for 51.1% of explained compositional variance. By contrast, neither anatomic site (*R*^2^ = 0.389; *P* = 0.876) nor traumatic etiology (*R*^2^ = 0.483; *P* = 0.626) influenced community composition (Table 3; Fig. 3). The large R^2^ values for non-significant comparisons most likely reflect unbalanced group sizes (Group A: n=123 vs Group B: n=44), which can inflate pseudo -F variance partitioning without generating statistical significance. PERMDISP (999 permutations) confirmed homogeneous within-group dispersions for the polymicrobial-versus-monomicrobial comparison (*F* = 2.85; *P* = 0.081) and the traumatic-versus-nontraumatic comparison (*F* = 0.28; *P* = 0.598), indicating that the significant PERMANOVA result for polymicrobial status reflects a genuine centroid shift rather than dispersion heterogeneity.

**FIG 3.**
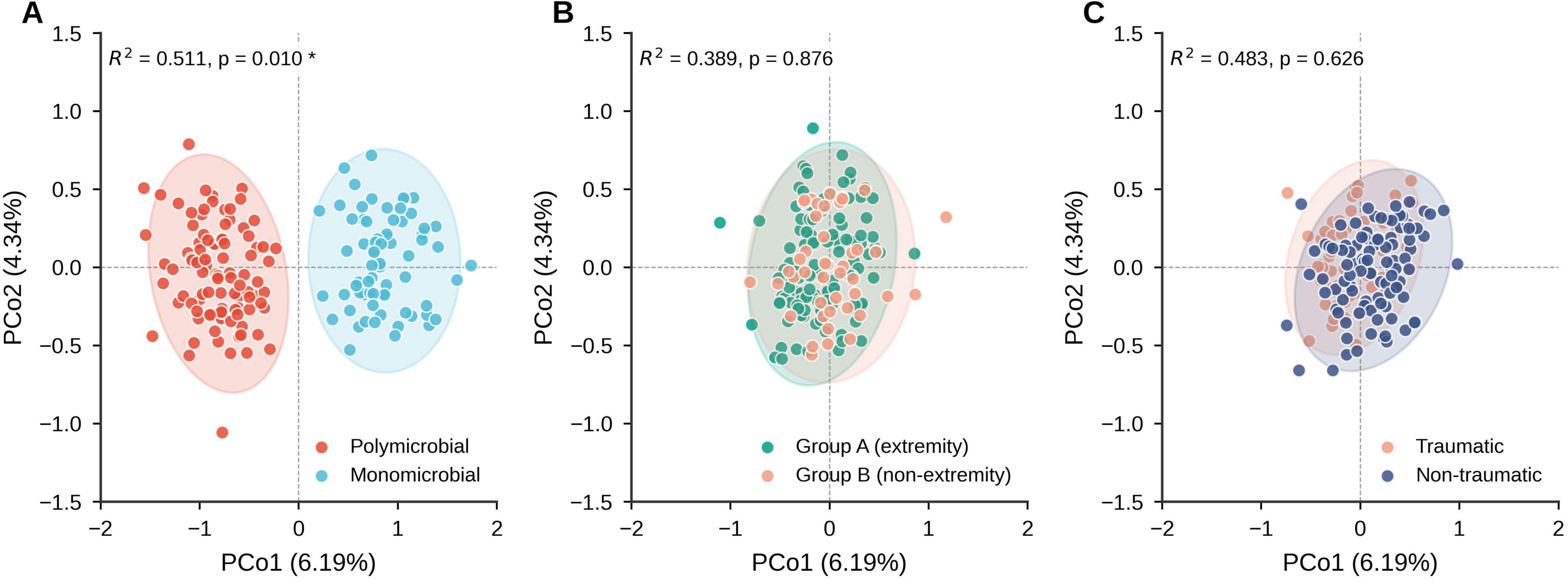
Principal-coordinate analysis (PCoA) of Bray-Curtis beta diversity across three clinical comparisons. Each point represents one patient tissue metagenome (n = 167). Shaded ellipses indicate 95% concentration ellipses. (A) Polymicrobial versus monomicrobial. (B) Group A extremity versus group B nonextremity. (C) Traumatic versus nontraumatic. PERMANOVA statistics (R² and P, 999 permutations) are annotated within each panel. PCo1 and PCo2 explain 6.19% and 4.34% of total variance, respectively.

**Table 3.** PERMANOVA and PERMDISP results: Bray-Curtis beta diversity (999 permutations).

| Comparison | n | PERMA<br>NOVA<br>Pseudo-F | PERMAN<br>OVA R<sup>2</sup> | PERMA<br>NOVA p | PERMD<br>ISP F | PER<br>MDIS<br>P p | Significance |
| --- | --- | --- | --- | --- | --- | --- | --- |
| Group A vs<br>Group B | 167 | 105.20 | 0.389 | 0.876 | , | , | ns |
| Traumatic vs<br>Non-traumatic | 167 | 154.22 | 0.483 | 0.626 | 0.28 | 0.598 | ns |
| Polymicrobial<br>vs<br>Monomicrobial | 163 | 85.55 | 0.511 | 0.010 | 2.85 | 0.081 | * (centroid<br>driven) |

PERMANOVA, permutational multivariate analysis of variance; PERMDISP, permutational test of multivariate homogeneity of group dispersions; R<sup>2</sup>, proportion of total community variance explained by grouping factor. PERMDISP was not computed for Group A vs Group B because no PERMANOVA significance was observed. The polymicrobial-versus-monomicrobial comparison excludes four culture-sterile samples (statistical n = 163). PERMDISP was performed on the first five PCoA axes (18.5% cumulative variance). ns, not significant; \* $p < 0.05$ .

### Co-occurrence network and differential abundance

Genus-level co-occurrence patterns were inferred from 16S abundance data. The resulting network revealed a tightly interconnected cluster of Gram-negative aerobes, *Acinetobacter*, *Escherichia*, *Klebsiella*, *Enterobacter*, and *Proteus*, linked by significant positive correlations across polymicrobial specimens (Fig. 4B). *Staphylococcus* and *Streptococcus* exhibited negative associations with this Gram-negative consortium, consistent with their predominance in monomicrobial infections. These patterns are consistent with synergistic interactions among enteric Gram-negative organisms, potentially mediated through proteolytic, hyaluronidase, and biofilm-related virulence determinants.

**FIG 4.**
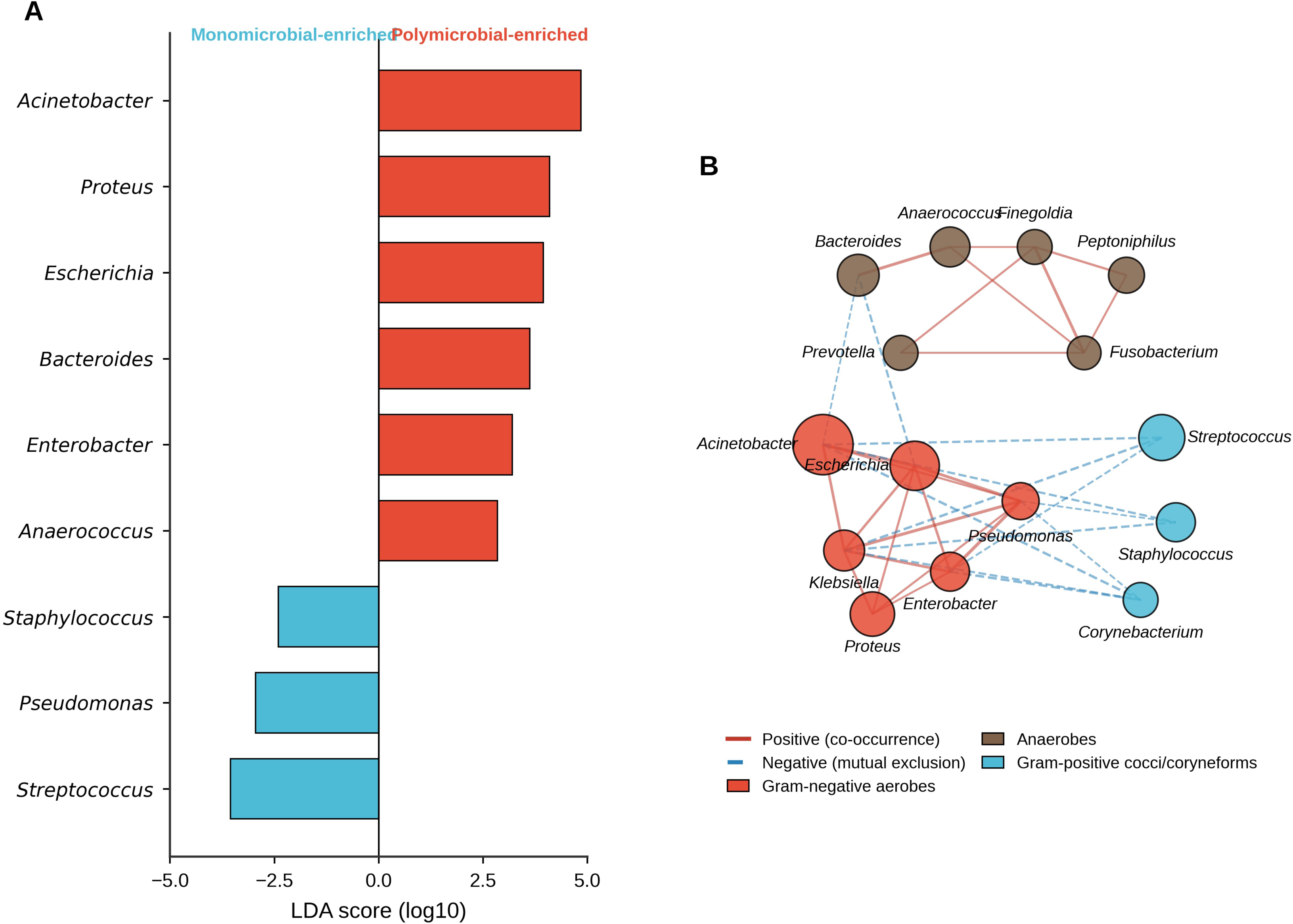
Differential abundance and microbial co-occurrence in polymicrobial versus monomicrobial NSTIs. (A) LEfSe horizontal bar plot of genera differentially abundant between polymicrobial and monomicrobial infections (LDA score, ≥2.0). (B) Co-occurrence network of the top abundant genera (n = 167); nodes represent genera (size proportional to mean relative abundance), and edges represent significant pairwise correlations (Spearman; SparCC permutations [n = 100]; adjusted P ≤ 0.05).

LEfSe analysis (LDA score, ≥2.0) identified genera differentially abundant between groups (Fig. 4A; see the supplemental material). *Acinetobacter* displayed the highest LDA score among polymicrobial-enriched genera (mean relative abundance, 18.1% versus 11.3% in monomicrobial infections), with *Proteus*, *Bacteroides*, *Escherichia*, and *Anaerococcus* also enriched in polymicrobial disease; *Streptococcus* and *Pseudomonas* were enriched in monomicrobial infections.

### Culture-metagenomics concordance and qPCR validation

Genus-level agreement between aerobic culture and 16S metagenomics was evaluated by both one-directional sensitivity and chance-corrected Cohen’s κ (Fig. 5C; see the supplemental material). The two metrics diverged substantially for several genera. *Escherichia* and *Klebsiella* showed almost perfect agreement (κ = 0.849 and 0.828; sensitivity, 88.9% and 80.0%), and *Proteus* showed substantial agreement (κ = 0.708; sensitivity, 71.0%). By contrast, the high one-directional concordance for *Bacteroides* (100.0%) and *Acinetobacter* (70.2%) corresponded to only fair κ values (0.327 and 0.289) once chance agreement was accounted for. *Streptococcus* (κ = 0.131) and *Enterococcus* (κ = 0.023) exhibited essentially no chance-corrected agreement, reflecting frequent metagenomics-positive/culture-negative discordance. Despite the modest κ for *Acinetobacter*, quantitative agreement at the load level was strong: qPCR *C*_T_ values correlated tightly with metagenomic relative abundance (Spearman ρ = −0.92; *P* < 0.0001; see the supplemental material).

**FIG 5.**
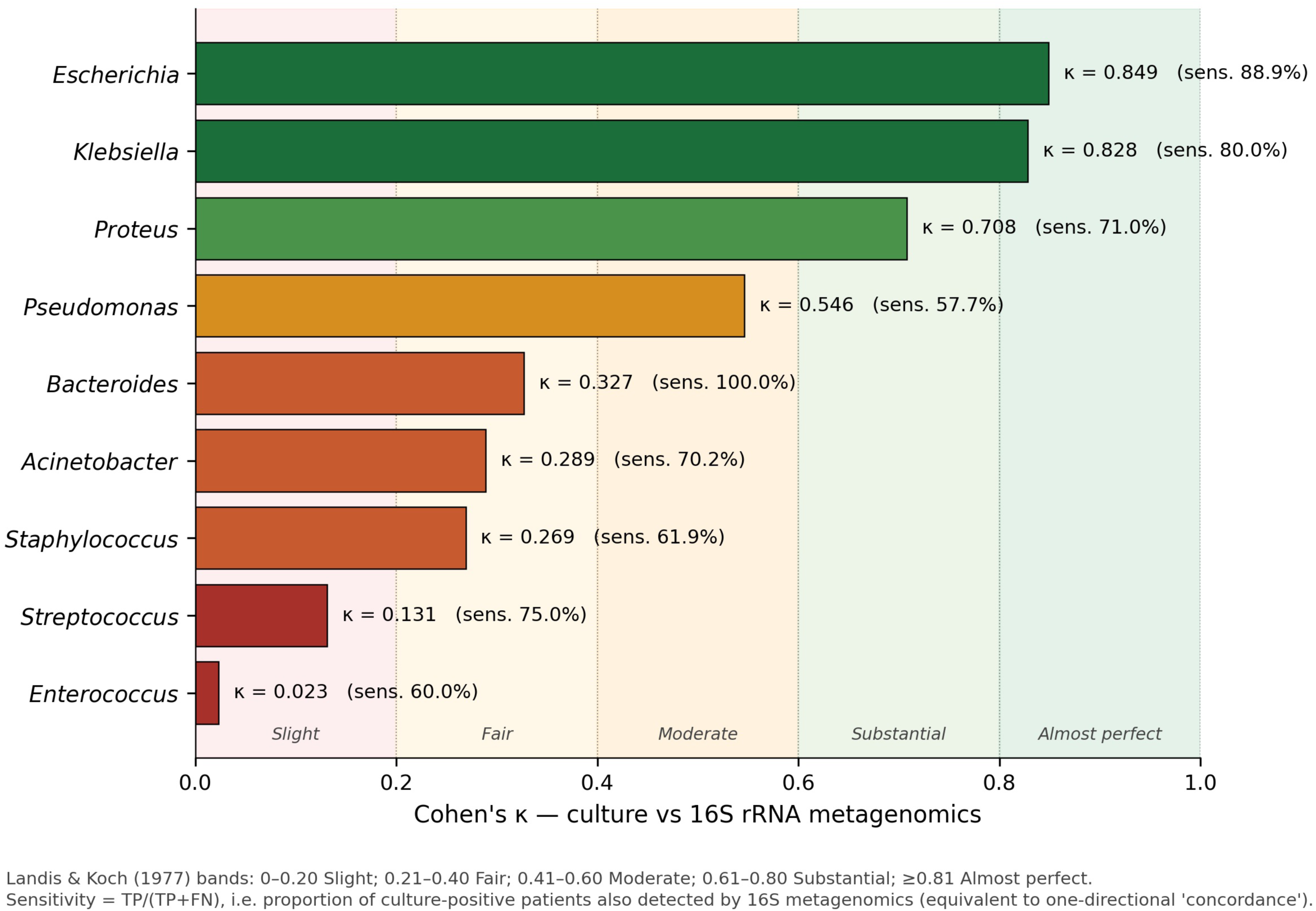
Three-way diagnostic comparison: culture, qPCR, and 16S metagenomics. (A) Detection rate of Acinetobacter baumannii by each modality. (B) Detection rate of Streptococcus pyogenes. Bars show patient-level positivity as a percentage of patients tested. (C) Genus-level culture-metagenomics agreement for nine taxa, plotted as Cohen’s κ on Landis-Koch bands (slight, 0 to 0.20; fair, 0.21 to 0.40; moderate, 0.41 to 0.60; substantial, 0.61 to 0.80; almost perfect, ≥0.81). One-directional sensitivity is annotated for comparison.

qPCR for *A. baumannii* was positive in 64/169 patients (37.9%), compared with 57/169 (33.7%) by culture and 82/167 (49.1%) by metagenomics (Fig. 5A). For *S. pyogenes*, qPCR was positive in 22/139 tested patients (15.8%), compared with 8/169 (4.7%) by culture and 51/167 (30.5%) by genus-level metagenomics (Fig. 5B). Thirty samples could not be tested by qPCR for *S. pyogenes* because of insufficient residual DNA following the *Acinetobacter* and 16S screening assays. The 3-fold qPCR-versus-culture gap and the 6-fold metagenomics-versus-culture gap together indicate substantial underdetection of *S. pyogenes* by conventional methods in this population, even where the genus contribution to total community burden remains modest.

## DISCUSSION

This study provides, to our knowledge, one of the first culture-independent characterizations of the NSTI wound microbiome from the Indian subcontinent. The dominant clinical signal is a Gram-negative, polymicrobial paradigm anchored by *A. baumannii* (33.7% by culture; 49.1% by metagenomics) and *E. coli* (32.0%), with *S. pyogenes* contributing only 4.7% of cases. Polymicrobial status, rather than anatomic site or traumatic etiology, was the principal determinant of community composition. The contrast with the Western evidence base is considerable: in the INFECT study, *S. pyogenes* accounted for 29 of 41 monomicrobial cases (5, 24), and polymicrobial infections were dominated by oral and gastrointestinal anaerobes (5, 12). The Gram-negative dominance we observe likely reflects tropical climate and soil contamination facilitating *A. baumannii* environmental inoculation through traumatic wounds, compounded by nosocomial acquisition during prolonged preoperative hospitalization (9–11, 28).

Alpha and beta diversity analyses collectively indicate that polymicrobial NSTIs represent a clinically and microbiologically distinct entity, not a quantitative extension of monomicrobial disease. A nonsignificant PERMDISP test (*F* = 2.85; *P* = 0.081) confirms that the PERMANOVA-detected separation reflects a genuine centroid shift in composition rather than heterogeneity of dispersion. Diversity metrics, particularly Shannon entropy, might therefore serve as culture-independent surrogates of polymicrobial status, a hypothesis warranting prospective validation.

The chance-corrected diagnostic comparison reveals a sharper picture than one-directional concordance alone permits. *Escherichia*, *Klebsiella*, and *Proteus* showed substantial to almost perfect agreement (κ ≥ 0.71), indicating that for these genera, the two modalities are largely interchangeable. By contrast, the apparently high concordance of *Bacteroides* (100% by sensitivity) and *Acinetobacter* (70.2% by sensitivity) shrinks to fair agreement once chance is accounted for (κ = 0.327 and 0.289), because metagenomics detects each genus in a large additional set of culture-negative samples. The very low κ for *Streptococcus* (0.131) and *Enterococcus* (0.023) similarly reflects a wide one-way detection gap. This pattern, high sensitivity coupled with low κ, is the molecular signature of fastidious, anaerobic, or nonviable organisms that aerobic culture systematically underrecovers (14, 15).

From an empirical-therapy perspective, in settings where *A. baumannii* dominates, broad-spectrum Gram-negative cover, typically a carbapenem-class agent paired with a glycopeptide for staphylococcal cover, should be prioritized, with antimicrobial stewardship adjustments as cultures finalize (1, 2, 25). The negligible role of *S. pyogenes* in this cohort indicates that intravenous immunoglobulin therapy is unlikely to yield benefit in this population, consistent with the negative INSTINCT trial (18). The Gram-negative co-occurrence cluster identified in our network analysis aligns with synergistic models in which enteric organisms cooperatively breach fascial planes through proteolytic, hyaluronidase, and biofilm-related virulence factors (22, 23). Because *A. baumannii* is a recognized biofilm former and the polymicrobial cluster includes additional biofilm-competent Enterobacterales, biofilm-mediated antimicrobial tolerance is a plausible contributor to high empirical-therapy failure rates in NSTI and merits direct investigation.

Strengths of this study include the large prospective cohort (*n* = 169), the multimodal diagnostic integration with chance-corrected concordance metrics, and the application of culture-independent methods to Indian NSTI. This study has several limitations. Amplicon-based 16S rRNA sequencing offers genus-level rather than species-level resolution, precluding direct attribution of antimicrobial resistance determinants. The cross-sectional design prevents temporal assessment of microbiome dynamics. The single-center nature, although partially mitigated by a wide geographic catchment area covering Punjab, Haryana, and Himachal Pradesh, might limit generalizability. The absence of shotgun metagenomics precludes functional annotation and resistome profiling. Extraction blanks and PCR no-template controls were not included in this sequencing run, a recognized concern for low-biomass interpretation; near-universal detection of *Enterococcus* and *Enterobacter* should therefore be interpreted with caution.

Longitudinal shotgun metagenomics integrated with dual host-pathogen RNA-seq, as pioneered by the INFECT investigators (5), would resolve virulence mechanisms within Gram-negative polymicrobial NSTIs. Quantitative point-of-care detection of *Acinetobacter* represents a pragmatic translational target for resource-limited settings. Multicenter Indian datasets are needed to confirm geographic generalizability and to inform region-specific empirical antibiotic guidelines. In summary, North Indian NSTIs present a microbial picture that is fundamentally distinct from the Western paradigm, with immediate implications for empirical antimicrobial therapy and for the design of region-specific clinical guidelines.

## ACKNOWLEDGMENTS

This study was funded by the Department of Science and Technology, Science and Engineering Research Board (DST-SERB; now the Anusandhan National Research Foundation), Government of India (grant EEQ/2021/000289). The funders had no role in study design, data collection, data analysis, data interpretation, or writing of the report. PGIMER, Chandigarh, provided fellowship support to the first author. We thank the Head of the Department of General Surgery, PGIMER, for assistance with intraoperative sample collection, the Department of Medical Microbiology, PGIMER, and all patients who consented to participate.

T.V. did the laboratory work, led the formal analysis, investigation, validation and drafted the original manuscript. N.R. and V.S. contributed to investigation, methodology, project administration and software. D.N. curated the data and methodology. and An.A. curated the data.

C.T. contributed to investigation through intraoperative sample collection and investigation. P.R. provided conceptualization, supervision and resources. Ar.A. and V.S. conceived and designed the study. Ar.A. supervised the project, conceptualization, acquired funding, and reviewed and edited the manuscript. N.R. and Ar.A. directly accessed and verified the underlying data and revised the draft. All authors read and approved the final manuscript.

## Declaration of generative AI use

During the preparation of this manuscript, the authors used Grammarly for language editing and refinement. After using this tool, the authors reviewed and edited the content as needed and take full responsibility for the content of the publication.

## Declaration of interests

We declare no competing interests.

## DATA AVAILABILITY

Raw 16S rRNA gene amplicon sequencing data have been deposited at the NCBI Sequence Read Archive under BioProject accession number PRJNA1466972. The data dictionary and deidentified analysis datasets will be available with publication. Patient-level metadata that could risk reidentification are not publicly available because of ethical and confidentiality restrictions but may be made available from the corresponding author on reasonable request and with approval of the Institutional Ethics Committee, PGIMER, Chandigarh, after a signed data access agreement, for noncommercial research. Custom Python analysis scripts are available from the corresponding author on reasonable request; all third-party software used is publicly available and cited in Materials and Methods.

**SUPPLEMENTAL MATERIAL** is available online.

---

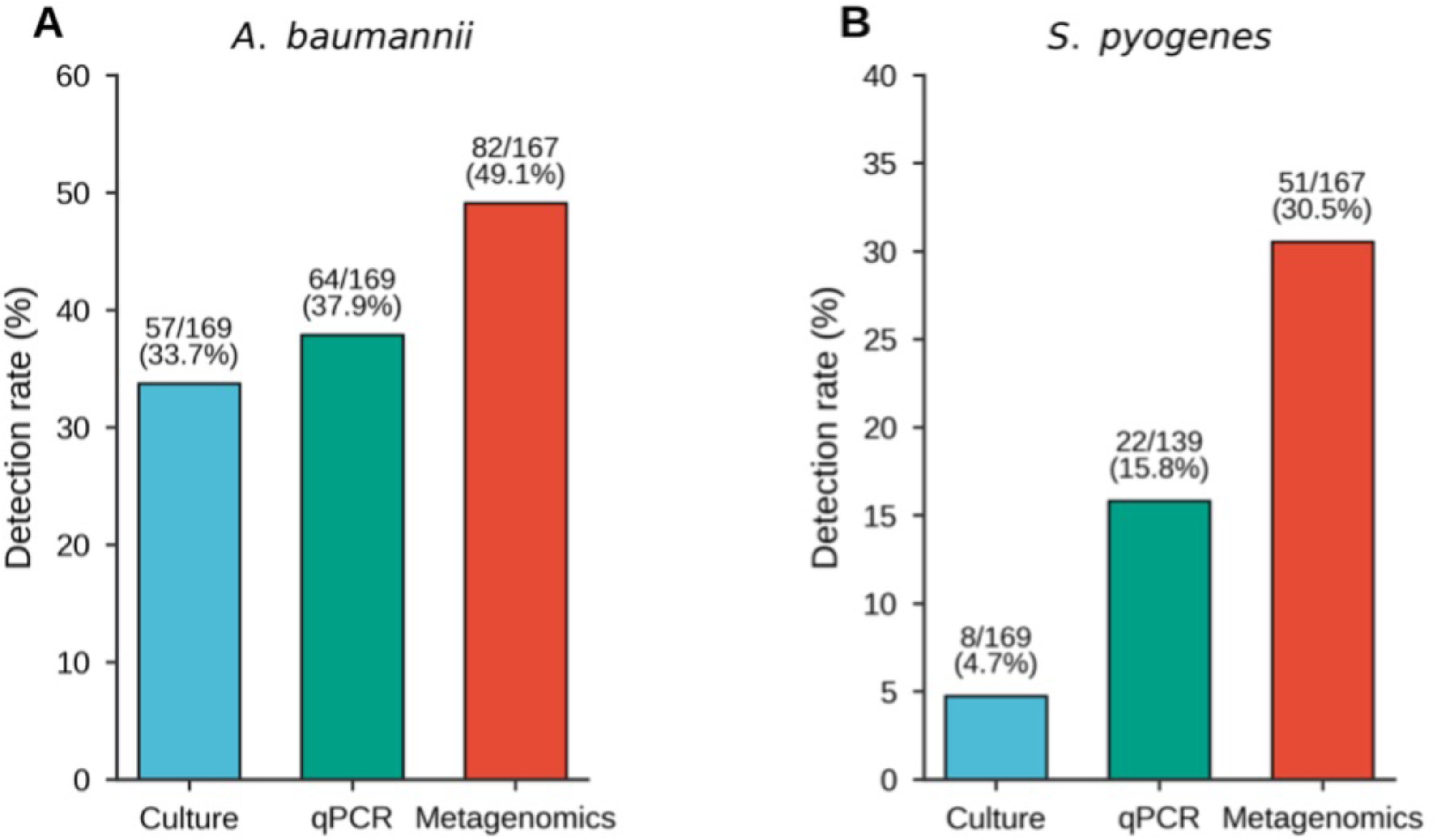

